# Traditional disease risk factors outperform epigenetic clocks as predictors of non-communicable disease morbidity in a middle-aged cohort

**DOI:** 10.64898/2026.02.10.705233

**Authors:** Daria Kostiniuk, Flóra Székely, Leo-Pekka Lyytikäinen, Jo Ciantar, Sonja Rajić, Pashupati P. Mishra, Terho Lehtimäki, Katja Pahkala, Suvi Rovio, Juha Mykkänen, Olli Raitakari, Emma Raitoharju, Saara Marttila

**Author notes:** These authors contributed equally to this work (Daria Kostiniuk, Flóra Székely).

## Abstract

DNA methylation-based epigenetic clocks have been highlighted as promising biomarkers of ageing, and they have been shown to robustly predict morbidity and mortality. However, current literature is lacking a formal analysis of the increased prediction accuracy, or the added value, of the epigenetic clocks over traditional risk factors of common chronic diseases.

Here, we have compared the most commonly used epigenetic clocks and traditional risk factors as predictors of incidence of ageing-associated non-communicable chronic disease in a 7-to-9-year follow-up in a middle-aged population cohort (n=1108, aged 34 to 49 years at baseline). In our cohort, a statistical model consisting of a combination of traditional risk factors outperforms any model including an epigenetic clock. The added value of epigenetic clock measurements over simple and affordable traditional risk factors should be clearly established, if epigenetic clocks are to be used in clinical settings or as tools of personal health monitoring.

## Main body

The concept of biological age aims to capture the amount of ageing-associated damage that has accumulated over time, and biomarkers of ageing are measures of this biological age. By definition, biomarkers of ageing should be better predictors of future health outcomes than mere chronological age (Baker & Sprott, 1988; Moqri et al., 2023). DNA methylation-based epigenetic clocks have been highlighted as promising biomarkers of ageing (Moqri et al., 2023, Duan et al., 2022). These epigenetic clocks have been shown to predict morbidity and mortality in population cohorts (Fransquet et al., 2019, Oblak et al., 2021, Chervova et al 2024). When the statistical models predicting morbidity or mortality are adjusted with common risk factors of non-communicable chronic diseases, for example BMI, smoking, alcohol consumption or exercise, the associations between epigenetic clocks and future health outcomes are usually retained, though they can be attenuated (Belsky et al., 2022; Lu et al., 2019; Hillary et al., 2020). Notably, traditional risk factors are also good predictors of morbidity and mortality on their own (Zhang et al., 2021; Barbaresko et al., 2018; Loef & Walach 2012; Chudasama et al., 2020). Unsurprisingly, these various risk factors associated with increased morbidity and mortality are also associated with increased epigenetic age (Wang et al., 2023; Nannini et al., 2022; Fox et al., 2023; Quach et al., 2017; Chervova et al., 2024, Oblak et al., 2021; Faul et al., 2023). As these risk factors are easy and affordable to measure, especially in comparison to DNA methylation, it is important to determine whether epigenetic clocks outperform them or provide additional predictive value by capturing aspects of biological aging not reflected by these simple risk factors. This is especially crucial if the epigenetic clocks are to be used in clinical settings or as tools of personal health monitoring.

Here, we have compared the predictive value of various epigenetic clocks and traditional risk factors of non-communicable chronic diseases in predicting disease incidence in a 7-to-9-year follow-up in the middle-aged Young Finns Study cohort (YFS, Pahkala et al., 2024). At baseline of this study setting, the participants were aged 34-49 years and were free from studied non-communicable chronic diseases, such as cardiometabolic diseases, hypertension, cancer and steatotic liver disease (Supplementary Table 1, Supplementary Table 2). Of the 1108 individuals included in the analysis, 222 (20.0%) were diagnosed with one or more non-communicable chronic disease or condition during the 7-to-9-year follow-up. Subjects with incident disease during the follow-up were older and had a higher BMI and waist-to-hip-ratio at baseline. The epigenetic clocks included in the analyses were age deviation (AgeDev) for Hannum (Hannum et al., 2013), Horvath (Horvath, 2013), PhenoAge (Levine et al., 2019) and GrimAge (Lu et al., 2019) and their principal component (PC) derivates (Higgins-Chen et al., 2023), as well as DunedinPACE (Belsky et al., 2022). For detailed description of methodology, see Supplementary Methods.

**Table 1.** Association between epigenetic clocks and incidence of any ageing-associated non-communicable chronic disease or condition (cardiometabolic diseases, hypertension, cancer, steatotic liver disease, Supplementary Table 2) in a 7-to-9-year follow-up (n=1108). Logistic regression models were adjusted for age, sex and Illumina array version. OR; odds ratio, 95% CI; 95% confidence interval.

|  | OR (95% CI) | p-value |
| --- | --- | --- |
| Horvath<sub>AgeDev</sub> | 1.00 (0.86 - 1.16) | 0.995 |
| PCHorvath<sub>AgeDev</sub> | 1.11 (0.95 - 1.29) | 0.181 |
| Hannum<sub>AgeDev</sub> | 1.08 (0.93 - 1.25) | 0.338 |
| PCHannum<sub>AgeDev</sub> | 1.07 (0.92 - 1.25) | 0.352 |
| PhenoAge<sub>AgeDev</sub> | 1.15 (0.99 - 1.34) | 0.066 |
| PCPhenoAge<sub>AgeDev</sub> | 1.18 (1.01 - 1.37) | **0.032** |
| GrimAge<sub>AgeDev</sub> | 1.20 (1.03 - 1.39) | **0.015** |
| PCGrimAge<sub>AgeDev</sub> | 1.13 (0.98 - 1.31) | 0.094 |
| DunedinPACE | 1.19 (1.02 - 1.39) | **0.024** |

In a minimally adjusted model (age, sex and Illumina array version), _PC_PhenoAge_AgeDev_, GrimAge_AgeDev_ and DunedinPACE were statistically significant predictors of non-communicable disease incidence during the 7-to-9-year follow-up (Table 1). These results are in line with published literature, as second- and third-generation clocks have been shown to be better predictors of future health outcomes as compared to first-generation clocks (Levine et al., 2018, Lu et al., 2019, Belsky et al., 2022).

For epigenetic clocks that were statistically significant predictors of non-communicable disease incidence in the minimally adjusted statistical model, we additionally adjusted the regression models with easy and affordable to measure risk factors of common chronic diseases, namely smoking, alcohol consumption, waist-to-hip-ratio (WHR) and BMI. In these fully adjusted models, none of the analysed epigenetic clocks were statistically significant predictors of disease incidence during follow-up (Supplementary Table 3).

We then compared a model consisting of the traditional risk factors (age, sex, smoking, alcohol consumption, WHR and BMI) to the minimally adjusted epigenetic clock models in relation to prediction of incidence of non-communicable diseases during the 7-to-9-year follow-up. The model consisting only of the traditional risk factors showed better discriminative performance as compared to any of the models containing the epigenetic clocks (Figure 1). However, none of the statistical models show particularly good discriminative performance. The differences in AUC between the traditional risk factor model and minimally adjusted epigenetic clock models were statistically significant (DeLong test p-value < 0.05, Supplementary Table 4).

**Figure 1.**
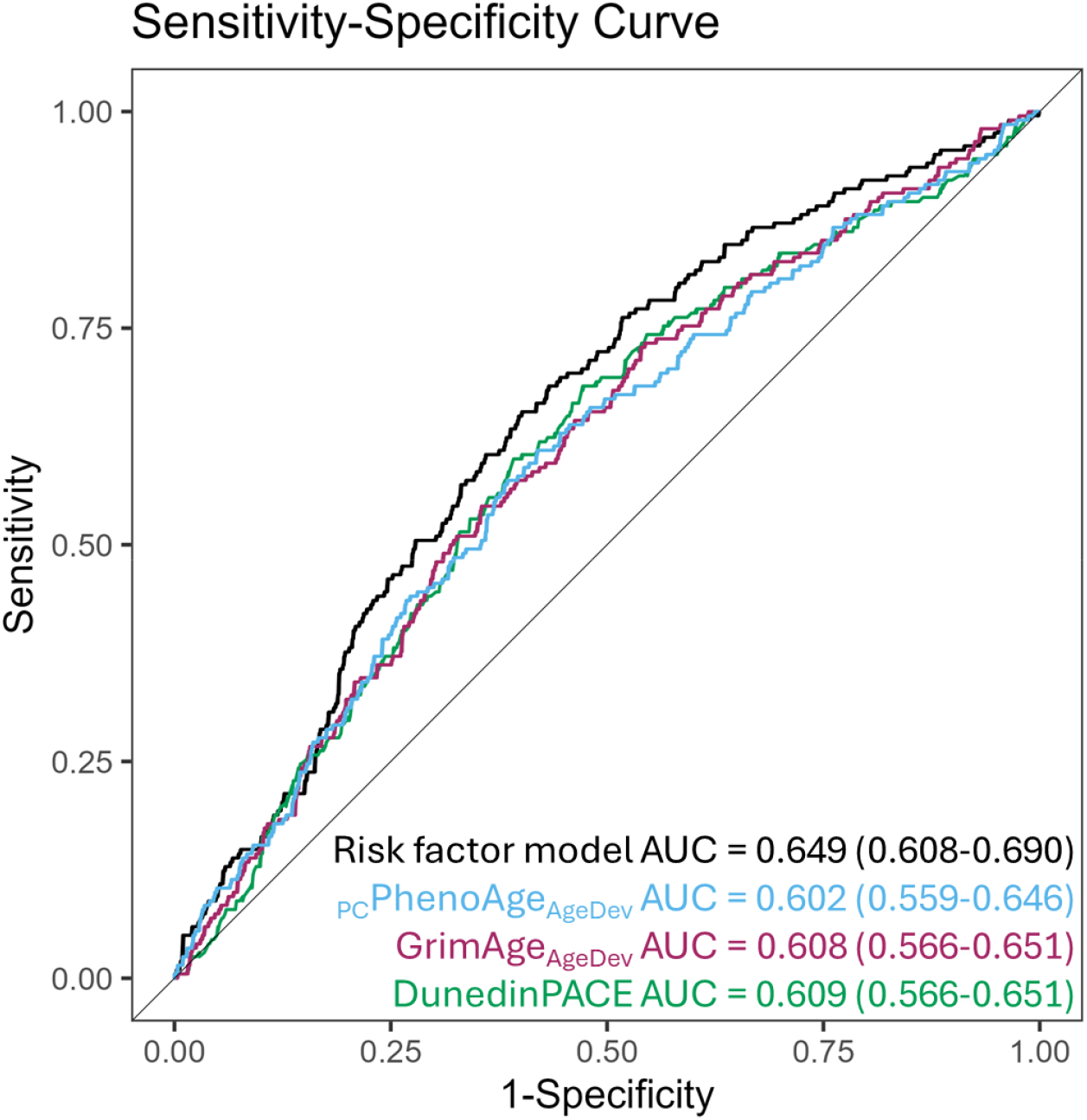
Receiver operating characteristics (ROC) curves comparing the discrimination performance of regression models consisting of traditional risk factors (risk factor model; age, sex, WHR, BMI, smoking, alcohol consumption) and minimally adjusted (age, sex, Illumina array) epigenetic clock models in relation to incidence of non-communicable disease during a 7-to-9-year follow-up.

In previous studies, adjusting for traditional risk factors has been shown to attenuate and even eliminate (Föhr et al., 2021) the association between epigenetic clocks and future health outcomes (Belsky et al., 2022; Lu et al., 2019; Hillary et al., 2020). Our finding that risk factors of common chronic disease outperform epigenetic clocks and the previously reported attenuation suggests that epigenetic clocks and traditional risk factors capture overlapping information, raising the question of whether these clocks contribute unique predictive value beyond traditional risk factors. The unique contribution of an epigenetic clock beyond established risk factors can be formally assessed using nested model comparisons, which test whether inclusion of the clock significantly improves model fit by adding information beyond one captured by traditional risk factors (Cook, 2018). In our study, such analyses were not feasible, as none of the studied clocks were statistically significant predictors of morbidity in fully adjusted models (Vickers et al., 2011). Notably, in studies where epigenetic clocks remain statistically significant predictors of future health outcomes after adjustments with traditional risk factors, formal assessments of their added predictive value have typically not been presented.

Compared to previous studies, where epigenetic clocks retained statistical significance also when adjusted for common risk factors, our cohort is younger and of a narrower age range, which could contribute to the poorer performance of epigenetic clocks in our cohort. Further studies in varying population cohorts, especially of varying age ranges, are needed to establish the demographic in which the epigenetic clocks would have the highest predictive performance and where they would provide the most added value. Especially interesting would be to establish whether the epigenetic clocks are predictive of future health outcomes in populations where the traditional risk factors are less informative, for example in secondary prevention, or in populations where there is less variation in the traditional risk factors, for example among individuals strictly adhering to healthy lifestyles.

As epigenetic clock measurements are already sold to consumers as tools of personal health monitoring, it should be shown they provide meaningful information also in the demographic most likely to purchase them. To the best of our knowledge, there is no data available on who are most likely to have their epigenetic age measured, but for other self-monitoring solutions, wearable devices, it is known that those who use in comparison to those who do not, are younger, healthier and more educated (Dhingra et al., 2023).

Epigenetic clocks potentially capture multiple aspects of the biological ageing process, and therefore when studying ageing itself or the molecular mechanisms of ageing, it is a more appropriate metric than a simple disease risk factor. However, if the aim is to predict future health outcomes for individuals, a simpler approach can be the cost-effective one. Given the higher cost of DNA methylation measurements, the added value of epigenetic clock measurements over traditional risk factors should be considerable to justify their use.

## Funding

The Young Finns Study has been financially supported by the following organisations: the Academy of Finland (grants 356405, 322098, 286284, 134309 (Eye), 126925, 121584, 124282, 129378 [Salve], 117797 [Gendi] and 141071 [Skidi]); the Social Insurance Institution of Finland; Competitive State Research Financing of the Expert Responsibility area of Kuopio, Tampere and Turku University Hospitals (grant X51001); the Juho Vainio Foundation; the Paavo Nurmi Foundation; the Finnish Foundation for Cardiovascular Research; the Finnish Cultural Foundation; the Sigrid Juselius Foundation; the Tampere Tuberculosis Foundation; the Emil Aaltonen Foundation; the Yrjö Jahnsson Foundation; the Signe and Ane Gyllenberg Foundation; the Diabetes Research Foundation of the Finnish Diabetes Association; EU Horizon 2020 (grant 755320 for TAXINOMISIS and grant 848146 for To Aition); the European Research Council (grant 742927 for MULTIEPIGEN project); the Tampere University Hospital Supporting Foundation; the Finnish Society of Clinical Chemistry; the Cancer Foundation Finland; pBETTER4U_EU (Preventing obesity through Biologically and bEhaviorally Tailored inTERventions for you; project number: 101080117); CVDLink (EU grant nro. 101137278); and the Jane and Aatos Erkko Foundation. Pashupati P. Mishra (grant no. 349708) and Emma Raitoharju (grants 330809 and 338395) were supported by the Academy of Finland.

Researchers in the current study have been supported by Yrjö Jahnsson Foundation, Juho Vainio Foundation, Päivikki and Sakari Sohlberg Foundation, Paulo Foundation, The Finnish Foundation for Cardiovascular Research and Pirkanmaa Regional Fund of the Finnish Cultural Foundation

## Availability of data and materials

The datasets generated and/or analysed during the current study are not publicly available due to restrictions imposed by Finnish legislation but are available from the corresponding author/data sharing committee upon a reasonable request.

## Author contributions

SM planned the study. SM and DK designed the data analysis. DK and FS performed the data analysis. L-PL, JC, SRa, PPM, TL, KP, SRo, JM and OR provided data. TL, OR, ER and SM aquired funding. SM wrote the first draft of the manuscript, DK participated in writing the first draft of the manuscript and all co-authors read and revised the final mansucript.

## Notes

### Competing Interest Statement

The authors have declared no competing interest.

