## Supplementary for "Traditional disease risk factors outperform epigenetic clocks as predictors of non-communicable disease morbidity in a middle-aged cohort"

**Supplementary Methods	2**

**Supplementary Tables	6**

**References	9**

**Supplementary Methods**

*Study population*

The Young Finns Study (YFS) is a multicenter follow-up study on cardiovascular risk from childhood to adulthood in Finland. The study was launched in 1980, with 3596 participants aged 3-18 years. The subjects were randomly selected from Finnish national registry among the chosen age groups and the five study districts (Raitakari et al., 2008; Pahkala et al., 2024). Data utilized in this study is from follow-ups of 2011 and 2018/2020. The examinations included physical measurements, blood tests, and questionnaires. The follow-up of 2011 included 2063 participants, who were aged 34-49 years at the time.

The study has been approved by the 1st ethical committee of the Hospital District of Southwest Finland on September 21st, 2010, and by local ethical committees (1st Ethical Committee of the Hospital District of Southwest Finland, Regional Ethics Committee of the Expert Responsibility area of Tampere University Hospital, Helsinki University Hospital Ethical Committee of Medicine, Research Ethics Committee of the Northern Savo Hospital District, and Ethics Committee of the Northern Ostrobothnia Hospital District). All study participants gave an informed consent, and the study was conducted according to the principles of the Declaration of Helsinki.

*Physical measurements and diseases*

Weight and height were measured, and BMI was calculated as weight(kg)/(height(m))^2^. Waist and hip circumference were measured twice, and the average was used to calculate waist-to-hip ratio (WHR). Blood pressure (BP) was obtained in-office using automatic BP monitor, measured in a sitting position with five-minute intervals between each of the three readings, and the average systolic and diastolic BPs were used for the analysis. Alcohol consumption and smoking habits were assessed using a questionnaire. Alcohol consumption was measured as drinks per day, and smoking was classified as daily smoker/non-smoker.

Information regarding diseases, medical conditions and medications (listed in Supplementary Table 2) were collected with a questionnaire, the participants were asked to report if they have any of the listed conditions diagnosed by a physician. Liver steatosis was evaluated from ultrasound imaging as described in Raitoharju et al., 2016, with individuals classified as having a clearly identifiable fatty liver considered as having a condition in our analyses. Individuals were considered to have hypertension if they indicated in the questionnaire that they had been diagnosed with hypertension or that they were taking hypertension medication or if their systolic BP was greater than 140 mmHg or diastolic BP greater than 90 mmHg. Individuals were considered to have type 2 diabetes if they indicated in the questionnaire that they had been diagnosed with type 2 diabetes or that they were taking oral glucose-lowering medication or insulin or if their fasting plasma glucose was ≥7.0 mmol/l or HbA1c ≥6.5.

*DNA methylation and epigenetic clocks*

DNA methylation analysis was performed from samples of the 2011 follow-up. Leukocyte DNA was obtained from EDTA blood using a Wizard® Genomic DNA Purification Kit (Promega Corporation, Madison, WI, USA) according to the manufacturer’s instructions. Genome-wide DNA methylation levels were obtained using Illumina Infinium BeadChips. 188 of the samples were analysed with Illumina Infinium HumanMethylation450 BeadChips as described previously in Kananen et al., 2016, and 1526 of the samples were analysed with Illumina Infinium MethylationEPIC BeadChips as described previously in Marttila et al., 2021. In total, for follow-up of 2011, methylation data was available for 1714 individuals. Array type was used as a covariate in the statistical analyses.

The epigenetic clocks utilized in the study were calculated according to published methods. The original clocks, Horvath as described in Horvath, 2013, Hannum as described in Hannum et al., 2013, PhenoAge as described in Levine et al., 2018 and GrimAge as described in Lu et al., 2019. DunedinPACE was calculated with the R package DunedinPACE as described in Belsky et al., 2022. The principal component derivates of the clocks were calculated as described in Higgins-Chen et al., 2022. All the clocks showed the expected strong correlation to chronological age (Supplementary Table 5). For all clocks utilised (except DunedinPACE) age deviation (AgeDev) was calculated by regressing the epigenetic age on chronological age (Chen et al., 2016; Moqri et al., 2023), and this AgeDev value was used in all statistical analyses.

*Statistical analyses*

Included in the analyses presented in this manuscript are individuals (n=1108) who had DNA methylation data available from 2011 follow-up and phenotypic data available from both 2011 and 2018/2020 follow-ups, and who were free of the included non-communicable chronic diseases, conditions and medications at baseline in 2011 (diseases, conditions and medications listed in Supplementary Table 2).

Association of the epigenetic clocks with the outcome (being diagnosed an incident disease or starting medication during the 7-to-9-year follow-up) was analysed with logistic regression. Minimally adjusted models were adjusted for age, sex and Illumina array version, and fully adjusted models included these same covariates in addition to BMI, WHR, smoking (daily smoker yes/no), and alcohol consumption (drinks per day). Predictive performance of logistic regression models in terms of discrimination was assessed using the area under receiver operating characteristic curve (AUROC). Differences in AUROC between non-nested models were assessed with two-sided DeLong test. 95% confidence intervals for AUROCs were estimated using DeLong’s method, with all calculations performed using the *pROC* package (Robin et al., 2011) and figures generated with the *ggplot2* package (Wickham 2016).

**Supplementary Tables**

**Supplementary Table 1.** Demographics of the study population at baseline (n=1108). At baseline, the study subjects did not have any of the conditions or medications listed in Supplementary Table 2.

|  |  | Remain healthy | Diseased in follow-up | p-value | Statistical test |
| --- | --- | --- | --- | --- | --- |
| n |  | 886 | 222 |  |  |
| Female (%) |  | 517 (58.4) | 124 (55.9) | 0.55 | chisq |
| Baseline age (mean (SD)) |  | 41.41 (4.99) | 43.07 (4.68) | <0.001 | t-test |
| Baseline BMI (median [IQR]) |  | 24.86 [22.48, 27.66] | 26.29 [23.74, 30.07] | <0.001 | u-test |
| Baseline waist-hip ratio (mean (SD)) |  | 0.88 (0.08) | 0.91 (0.08) | <0.001 | t-test |
| Daily smoker at baseline (%) | Yes | 115 (13.4) | 25 (12.0) | 0.669 | chisq |
| Baseline drinks per day continuous (median [IQR]) |  | 0.43 [0.14, 0.86] | 0.43 [0.14, 0.86] | 0.647 | u-test |
| Baseline drinks per day categorical (%) | 0 | 193 (22.8) | 44 (21.8) | 0.913 | chisq |
|  | 0-2 | 571 (67.6) | 136 (67.3) |  |  |
|  | 2-4 | 69 (8.2) | 18 (8.9) |  |  |
|  | >4 | 12 (1.4) | 4 (2.0) |  |  |

**Supplementary Table 2.** Incident non-communicable disease frequencies at follow-up. One individual could gain more than one condition; therefore, the sum of frequencies exceeds the total number of individuals who gained a condition during follow-up (n=222).

| **Disease frequencies at follow-up** | **Frequency** |
| --- | --- |
| Hypertension* | 184 |
| Other arrhythmia | 66 |
| Cholesterol medication | 64 |
| Cancer | 28 |
| Type 2 diabetes** | 24 |
| Valvular disease | 22 |
| Liver steatosis*** | 17 |
| Coronary heart disease | 15 |
| Coronary artery angioplasty | 12 |
| Transient ischemic attack | 10 |
| Myocardial infarction | 9 |
| Osteoporosis | 9 |
| Atrial fibrillation | 6 |
| Carotid artery stenosis | 6 |
| Ischemic stroke | 4 |
| Heart failure | 3 |
| Claudication | 3 |
| Dilation of aorta | 2 |
| Haemorrhagic stroke | 2 |
| Bypass surgery of coronary arteries | 2 |
| Peripheral artery angioplasty | 2 |
| *Based on self-reported diagnosis or antihypertensive medication | |
| ** Fasting plasma glucose ≥7.0 mmol/l, or HbA1c ≥6.5, or use of oral glucose-lowering medication or insulin (but not type 1 diabetes) or those who had been diagnosed with T2D by a physician. | |
| ***As measured by ultrasound | |

**Supplementary Table 3**. Association between epigenetic clocks that were significant in minimally adjusted model (Table 1) and incidence of any ageing-associated non-communicable chronic disease or condition (cardiometabolic diseases, hypertension, cancer, steatotic liver disease, for detailed list see Supplementary Table 2) in a 7-to-9-year follow-up (n=1039) in the fully adjusted logistic regression model (age, sex, Illumina array version, smoking, alcohol consumption, WHR, BMI). OR; odds ratio, 95% CI; 95% confidence interval.

|  | OR (95% CI) | p-value |
| --- | --- | --- |
| _PC_PhenoAge_AgeDev_ | 1.12 (0.96 - 1.32) | 0.156 |
| GrimAge_AgeDev_ | 1.20 (0.98 - 1.47) | 0.076 |
| DunedinPACE | 1.05 (0.88 - 1.26) | 0.563 |

***Supplementary Table 4.*** *Comparison of the discrimination performance between minimally adjusted (age, sex, Illumina array) epigenetic clock models and a model composed of easy and affordable to measure risk factors (age, sex, smoking, alcohol consumption, WHR, BMI) models in relation to incidence of non-communicable disease during a 7-to-9-year follow-up. Higher AUC values indicate better discriminatory performance. Differences in AUC between models were assessed with DeLong test. For the ROC curves, see Figure 1.*

|  | n | AUC | DeLong test p-value |
| --- | --- | --- | --- |
| Risk factors | 1039 | 0.649 (0.608-0.690) | NA |
| _PC_PhenoAge_AgeDev_ | 1039 | 0.602 (0.559-0.646) | 0.02132 |
| GrimAge_AgeDev_ | 1039 | 0.608 (0.566-0.651) | 0.03466 |
| DunedinPACE | 1039 | 0.609 (0.566-0.651) | 0.02616 |

**Supplementary Table 5.** Correlation between chronological age and the different epigenetic clocks used in the study (n=1108). Note, that while age deviation (AgeDev) values were used in regression analyses presented in the main body and in the supplementary material, these correlations are presented for the epigenetic clock values themselves (representing epigenetic age in years).

|  | Spearman r | p-value |
| --- | --- | --- |
| Horvath | 0.700 | <1*10^-22^ |
| _PC_Horvath | 0.728 | <1*10^-22^ |
| Hannum | 0.714 | <1*10^-22^ |
| _PC_Hannum | 0.732 | <1*10^-22^ |
| PhenoAge | 0.704 | <1*10^-22^ |
| _PC_PhenoAge | 0.648 | <1*10^-22^ |
| GrimAge | 0.731 | <1*10^-22^ |
| _PC_GrimAge | 0.824 | <1*10^-22^ |
| DunedinPACE | 0.162 | 5.92*10^-8^ |
